# A novel LysinB from an F2 sub-cluster mycobacteriophage *RitSun*

**DOI:** 10.1101/2024.02.29.582697

**Authors:** Ritu Arora, Kanika Nadar, Urmi Bajpai

**Author notes:** **Address for correspondence:** Urmi Bajpai, PhD (Professor), Department of Biomedical Science, Acharya Narendra Dev College (University of Delhi), Govindpuri, Kalkaji, New-Delhi-110019, India., Email id.

## Abstract

With the growing antibiotic resistance in mycobacterial species posing a significant threat globally, there is an urgent need to find alternative solutions. Bacteriophage-derived endolysins aid in releasing phage progeny from the host bacteria by attacking the cell wall at the end of their life cycle. Endolysins are attractive antibacterial candidates due to their rapid lytic action, specificity and low risk of resistance development. In mycobacteria, owing to the complex, hydrophobic cell wall, mycobacteriophages usually synthesize two endolysins: LysinA, which hydrolyzes peptidoglycan; LysinB, which delinks mycolylarabinogalactan from peptidoglycan and releases mycolic acid. In this study, we conducted domain analysis and functional characterization of a recombinant LysinB from *RitSun*, an F2 sub-cluster mycobacteriophage. Several properties of *RitSun* LysinB are important as an antimycobacterial agent: its ability to lyse *Mycobacterium smegmatis* ‘from without’, a specific activity of 1.36 U/mg, higher than the reported ones and its inhibitory effect on biofilm formation. Given the impervious nature of the mycobacterial cell envelope, native endolysins’ ability to damage cells on exogenous applications warrants further investigation. A molecular dissection of *RitSun* LysinB to identify its cell wall destabilizing sequence could be utilized to engineer other native lysins as fusion proteins and expand their activity profile.

## INTRODUCTION

The Mycobacterium genus consists of over 150 species, many of which are human pathogens. Tuberculosis (TB) caused by *Mycobacterium tuberculosis* (Mtb) is the world’s leading cause of death from a single infectious disease, with an estimated 1.6 million deaths in 2021 [1]. The estimated number of deaths has increased between 2019 and 2021, reversing many years of progress in reducing the TB burden. Antibiotic resistance is a widely acknowledged problem, identified by WHO as one of the most severe threats to global health, food security, and development today and has predicted it will inevitably result in higher medical costs, extended hospital stays, and higher mortality rates [2,3]. This chain of events, in turn, could damage any country’s economic, political, and social aspects. This has geared the scientific community to increase their efforts toward exploring alternatives to standard antibiotic therapy. Though TB is curable, the emergence of multi-drug-resistant (MDR) Mtb strains has severely thwarted the progress toward eliminating TB. With the global burden of MDR-TB increasing by more than 20% per year over the past several years, drug-resistant TB is expected to kill 75 million people over the next 35 years, emphasizing the need to develop new alternative anti-tuberculosis therapies [4]. The infections caused by non-tuberculous mycobacteria (NTM) also threaten animals and human health [5] and have emerged as a new threat, especially to the immunocompromised population. Hence, there is an urgent need to find new drugs and alternative solutions [6].

Bacteriophages, commonly known as phages, are viruses that infect and replicate in bacterial hosts and are being explored as alternatives to antibiotics with renewed vigour. Phages are ubiquitous and are estimated as the most abundant (10^31^) biological entity on earth [7], and their encoded lytic proteins, such as endolysins, are also emerging as non-antibiotic options to treat drug-resistant bacterial infections. Mycobacteriophages are phages that infect mycobacterial hosts, including members from the *M. tuberculosis* complex and non-tubercular mycobacterial (NTM) hosts.

Most double-stranded DNA (dsDNA) lytic bacteriophages end the infection cycle through programmed and controlled actions of a lytic cassette, which includes holin and endolysins. Holin is a membrane protein that oligomerizes and forms a channel in the bacterial cell membrane for endolysins to access the peptidoglycan, which targets the peptidoglycan and disintegrates it for new phage progeny to release [8,9]. Unlike antibiotics, endolysins are specific to the target bacteria and hence do not harm the normal commensal microflora, are effective against drug-resistant strains of pathogens and offer additional advantages in targeting both growing and non-growing bacterial cells. They can either be used in native form or can be engineered to enhance their properties, desired in clinical applications or as biocontrol agents.

Endolysins emerged as promising anti-Gram-positive bacterial agents since they could lyse the bacteria due to direct contact with the peptidoglycan [10,11,12,13]. However, in the case of Gram-negative bacteria and mycobacterial spp., the outer membrane/envelope acts as a barrier for enzymes to access the peptidoglycan [14,15]. Hence, interventions such as outer membrane permeabilizers are required [16,17,18,19].

The first reports on the use of endolysins to treat drug-resistant infections came in the early years of this century [20]. Exebacase (lysin CF-301), which targets *Streptococcus* and *Staphylococcus* species to treat infective endocarditis, was the first to enter human clinical trials in the United States and succeeded in phase I and II [21,22]. Endolysin Sal200 targets antibiotic-resistant *Staphylococcus* infections and has been successful in phase I clinical trials [21,23]. Lysins have been successfully demonstrated as commercial products; an example is Gladskin, developed by Micreos, which targets *S. aureus* infections, causing skin disorders such as eczema and acne [21,24].

The prevalence of drug-resistant strains of *M. tuberculosis* and a rise in infections caused by non-tuberculous mycobacterial (NTM) pathogens demand an expanding repertoire of lysins, which can be screened for their lytic efficiency, structural diversity and host-spectrum. However, fewer reports are available on lysins against Mycobacterial spp. The role of mycobacteriophages and their derived endolysins in human therapy has gained attention, especially in the last few years when mycobacteriophages were used to treat drug-resistant NTM infections [25,26]. Phages and endolysins can also play an important role in potentially shortening the treatment regimen of TB/NTM infections [9,27]. The mycobacterial cell envelope is more elaborate when compared to Gram-positive and Gram-negative bacteria; it contains an additional mycolylarabinogalactan layer, which forms an ester linkage with the peptidoglycan. Due to the complexity of the cell wall, mycobacteriophages encode two endolysins: LysinA and LysinB. At the end of the phage life cycle, holin helps in translocating endolysins to the periplasm, where LysinA acts upon the peptidoglycan, and LysinB cleaves the ester linkage between mycolic acid and mycolylarabinogalactan and also degrades the mycolic acid.

LysinB enzymes are unique to mycobacteriophages and have been examined more than LysinA. Exogenous administration of purified LysinB enzyme from *D29* has shown lysis from without and disrupted mycobacterial biofilms [14,28]. Mycobacteriophage endolysins have also been demonstrated to be utilised on functionalized nanobeads to protect surfaces such as personal protective equipment (PPE) that routinely comes into contact with aerosolised bacteria [28].

Furthermore, the recent reports on *D29* [29,30] and *Ms6* Lysin B [31] have provided promising evidence supporting LysinB as a new therapeutic strategy.

In this report, we present the purification and characterization of a novel LysinB from an F2 cluster mycobacteriophage ‘*RitSun*’ isolated on *M. smegmatis* Mc^2^ 155. We studied its domain organization using online computational tools and performed sequence comparison with LysinB from the other F sub-clusters. We tested the esterase activity of the recombinant LysinB using an *in vitro* biochemical assay, and to assess the inhibitory effect of *RitSun* LysinB on mycobacterial growth, assays were performed on planktonic cells and the biofilm. With the planktonic cells, plate lysis assay and microtiter plate-based turbidity reduction method and the inhibitory effect on biofilm was carried out using Crystal Violet staining. The morphology of the enzyme-treated *M. smegmatis* Mc^2^ 155 cells was examined by FESEM analysis.

## RESULTS AND DISCUSSION

### 1. *In silico* characterization of *RitSun* LysinB

#### 1.1 Phylogenetic Analysis of *RitSun* LysinB gene

Phylogenetic analysis of *RitSun* LysinB gene sequence was performed against a database of double-stranded DNA viral sequences of different families and hosts (Figure 1A). As of today, the F2 sub-cluster has eight mycobacteriophages (including *RitSun)*. Expectedly, we found the other seven F2 sub-cluster mycobacteriophages (marked with a red circle: Figure 1B) as the close relative of *RitSun* LysinB and *Che9d* as the closest homolog. *RitSun* LysinB sequence also showed similarities with phages belonging to different F sub-clusters (Figure 1B).

**Figure 1.**
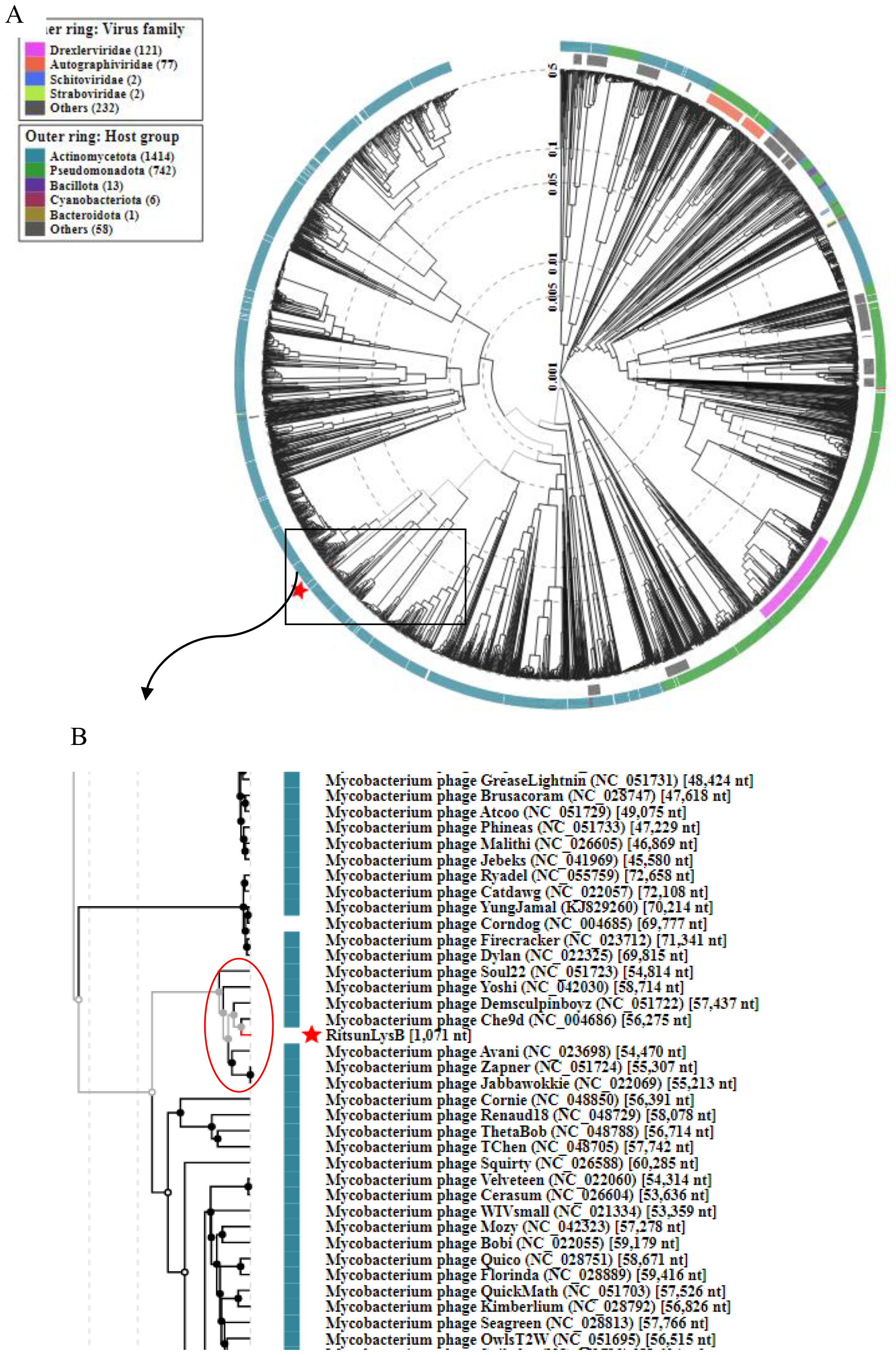
Phylogenetic analysis of *RitSun* LysinB gene sequence using VipTree. The reference list has 2300 sequences belonging to double-stranded prokaryotic DNA viruses from various families. **Figures 1A** and **1B** represent the circular phylogeny tree and a small portion of the rectangular phylogeny tree, respectively. All eight F2 sub-cluster mycobacteriophages are marked with a red circle.

#### 1.2 Domain Organization and Sequence Alignment

*RitSun* LysinB is a serine esterase based on the predicted conserved motifs and domains. InterProScan analysis predicted the presence of the characteristic conserved motifs: G-X-S-X-G pentapeptide (here, X_1_=F and X_2_=R) and GXP (X=N), a highly conserved motif found in lipases [32]. The predicted domains are alpha/beta hydrolase fold and a C-terminal linker domain (Figure 2A).

**Figure 2A.**
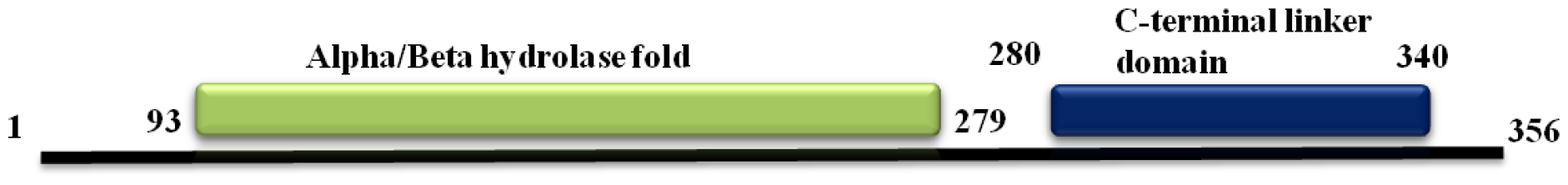
InterProScan-predicted domain organization of *RitSun* LysinB.

**Figure 2B.**
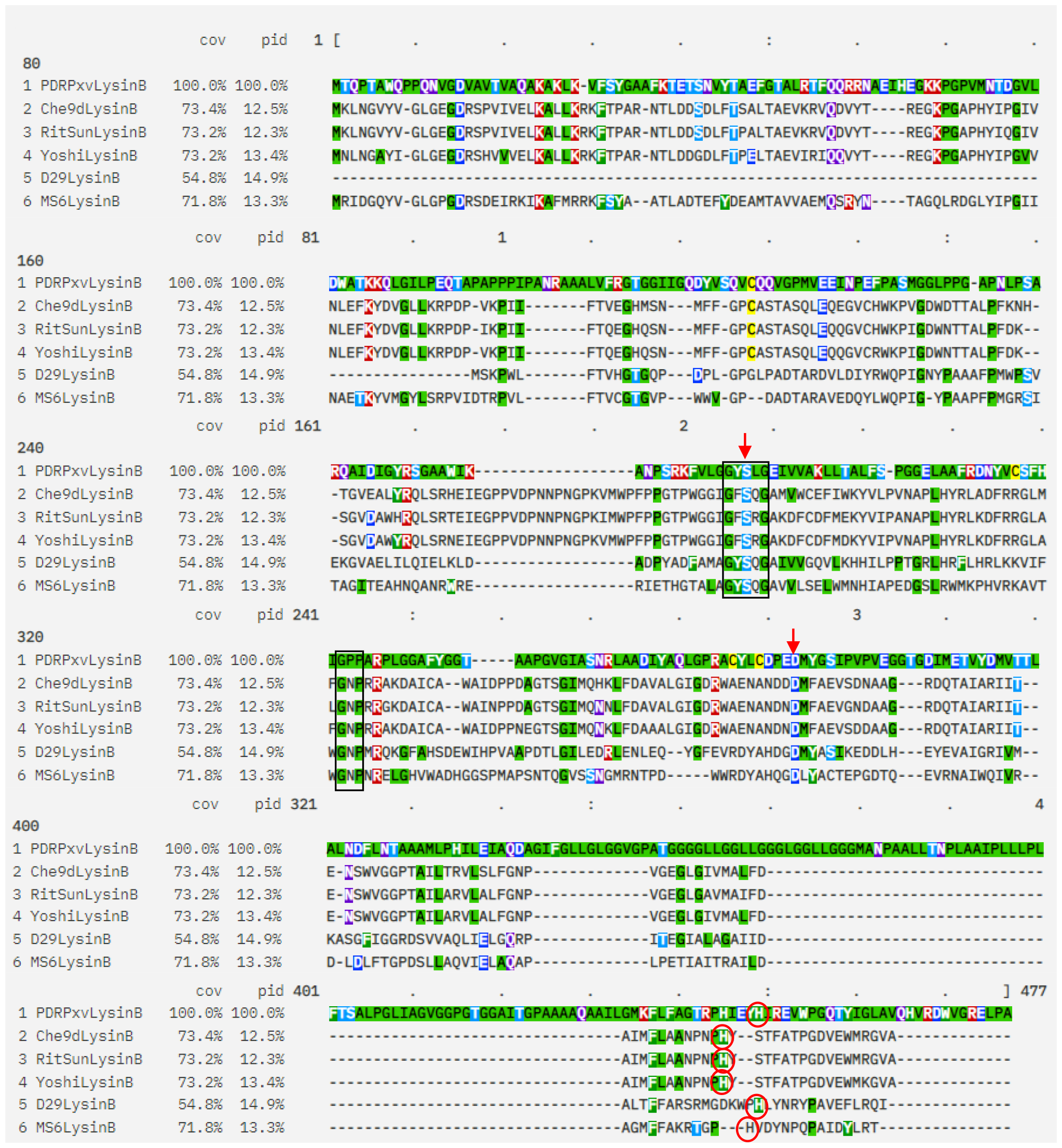
Sequence Alignment of LysinB from *RitSun, Yoshi, Che9d, Ms6, D29*, and *PDRPxv* mycobacteriophages using Clustal Omega Analysis. The results were observed using Mview. The conserved pentapeptide GXSXG and GXP motifs are marked with a square box. Ser and Asp of the catalytic triad are present at positions 186 and 270, respectively and marked with a red arrow aligned with that of D29 LysinB, whereas His (339) is shifted by two positions towards N terminal in *RitSun* LysinB.

**Figure 2C.**
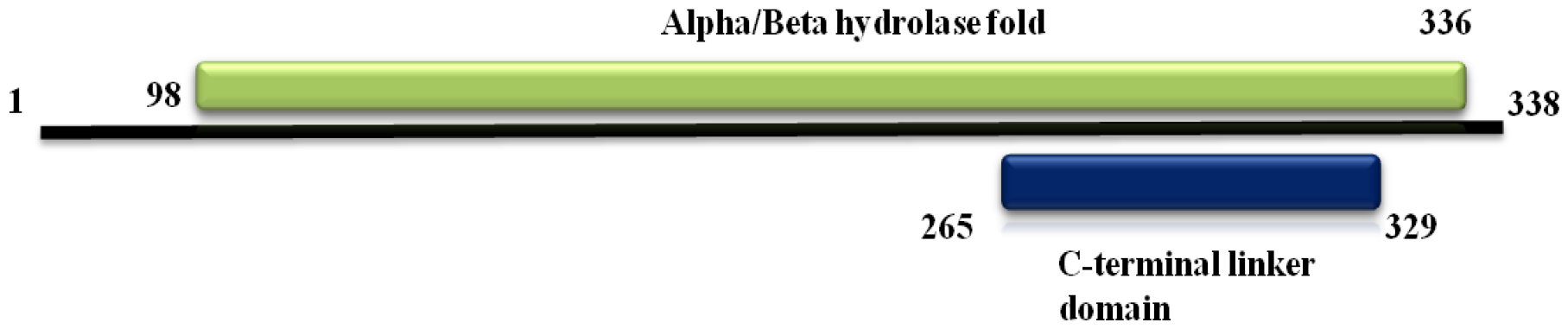
InterProScan-predicted domain organization of *Flathead* LysinB, which shows 28.86% identity (query cover of 95%) with *RitSun* LysinB.

LysinB enzymes, reported from other mycobacteriophages, also contain alpha/beta hydrolase fold and a C-terminal linker domain. In addition to these domains, *D29* LysinB has a Cutinase/Acetyl xylan esterase domain embedded in the alpha/beta hydrolase fold (Table 1).

**Table 1.** Characteristic properties of LysinB from different mycobacteriophages.

| LysinB | No. of amino acids | Domain(s) | Specific Activity (U/mg) | Reference |
| --- | --- | --- | --- | --- |
| *RitSun* | 357 | Alpha/beta hydrolase fold, C-linker domain | 1.36 | This study |
| *PDRP<sub>xv</sub>* | 451 | Alpha/beta hydrolase fold, A glycine rich C-terminal | 0.56 | [13] |
| *D29* | 254 | Alpha/beta<br>hydrolase fold,<br>Cutinase/acetyl<br>xylan esterase<br>domain, C-linker<br>domain | 0.72 | [14] |
| *Ms6* | 332 | Alpha/beta<br>hydrolase fold, C-linker domain | 0.12 | [33] |
| *Yoshi* | 356 | Alpha/beta<br>hydrolase fold, C-linker domain | - | Unpublished,<br>Sequence<br>obtained from<br>phagesdb.org<br>[34] |
| *Che9d* | 357 | Alpha/beta<br>hydrolase fold, C-linker domain | - | Unpublished,<br>Sequence<br>obtained from<br>phagesdb.org<br>[34] |

The enzymatic activity of *D29* LysinB has been demonstrated to depend on the presence of the catalytic triad Ser82-Asp166-His240, with serine being a constituent of the pentapeptide. *Fraga et al*. [29] reported Ser82-Ala82 mutation to abolish its activity, indicating serine as a key residue for the enzymatic activity. Also, while Ser82 and Asp166 are highly conserved residues in the catalytic triad, His shows a shift either by one position towards the N-terminal or three positions towards the C-terminal [35,13]. In Figure 2B, we have demonstrated an alignment of *RitSun* LysinB with *Ms6* LysinB (F1 sub-cluster), *D29* LysinB (A2 sub-cluster), *PDRPxv* LysinB (B1 sub-cluster), *Che9d* LysinB (F2 sub-cluster) and *Yoshi* LysinB (F2 sub-cluster). *Yoshi* LysinB shows the highest identity with *RitSun* Lysin B, and *Che9d* shares the same clade and is observed to originate from a common ancestor; hence, they are taken here as representative gene sequences from the F2 sub-cluster. In *RitSun, Che9d* and *Yoshi* LysinB, His residue is shifted by two positions towards the N-terminal (marked with a circle); *Ms6* LysinB has a shift of single position, whereas *PDRPxv and D29* LysinB do not show a shift in His residue and shows an alignment with each other.

#### 1.3 Comparative analysis of *RitSun* LysinB with phages from other F sub-clusters

The F cluster is divided into five sub-clusters: F1 to F5, and the F2 sub-cluster has seven mycobacteriophages reported to date (excluding *RitSun*) [34]. We used Blastp to compare the *RitSun* LysinB sequence with LysinB from the other F sub-clusters phages. Out of seven sub-cluster phages, *RitSun* LysinB revealed maximum identity (91.29%) with LysinB from mycobacteriophage *Yoshi*, and a minimum identity (83.71%) with LysinB from mycobacteriophage *Soul22*.

F1 is the most populated sub-cluster, with 221 members. We extracted the LysinB protein sequence from 150 F1 sub-cluster phage for comparative analysis. We found *RitSun* LysinB to show >83% identity (100% query cover) with seven F1 phages: *EleanorGeorge, Enby, Girr, Kingsley, Lorde, MisterCuddles and Ovechkin* and 84.73% identity, with a query cover of 97%, with *Hamulus* LysinB. All the other phages from the F1 sub-cluster showed less than 30% identity when the query cover was ≥ 95%. The F3 sub-cluster has only one phage reported, and its genome, a Lysin gene, has not been predicted [34]. The F4 sub-cluster has four phages, out of which LysinB from phages *ThetaBob* and *TChen* showed 88.52% and 88.24% sequence identity at 100% query cover, respectively. The F5 sub-cluster has only one reported phage. Its LysinB showed 28.65% identity (with a 99% query cover) with *RitSun* LysinB.

Using InterProScan, we also performed a comparative domain analysis of *RitSun* LysinB with LysinB from other F sub-cluster phages. LysinB protein sequences from F2, F4, and F5 sub-cluster phages and *EleanorGeorge, Hamulus*, and *Flathead* phages from the F1 sub-cluster were compared. We observed that LysinB sequences showing a high identity with *RitSun* LysinB (>83%) show a similar architecture: an α/β hydrolase fold followed by a linker domain (Figure 2A). Interestingly, those showing lower sequence identity (<30%) were predicted to have a C-terminal linker domain embedded within the α/β hydrolase fold. The domain organisation of a low identity (28.86%) LysinB from *Flathead* phage is shown in Figure 2C.

### 2. Purification of *RitSun* LysinB and Western Blot Analysis

The His-tagged *RitSun* LysinB was overexpressed in *E. coli* BL21 (DE3), and the protein was purified by Ni-NTA affinity chromatography (Figure 3A). Western blotting using anti-His antibodies confirmed the presence of purified recombinant LysinB (Figure 3B).

**Figure 3.**
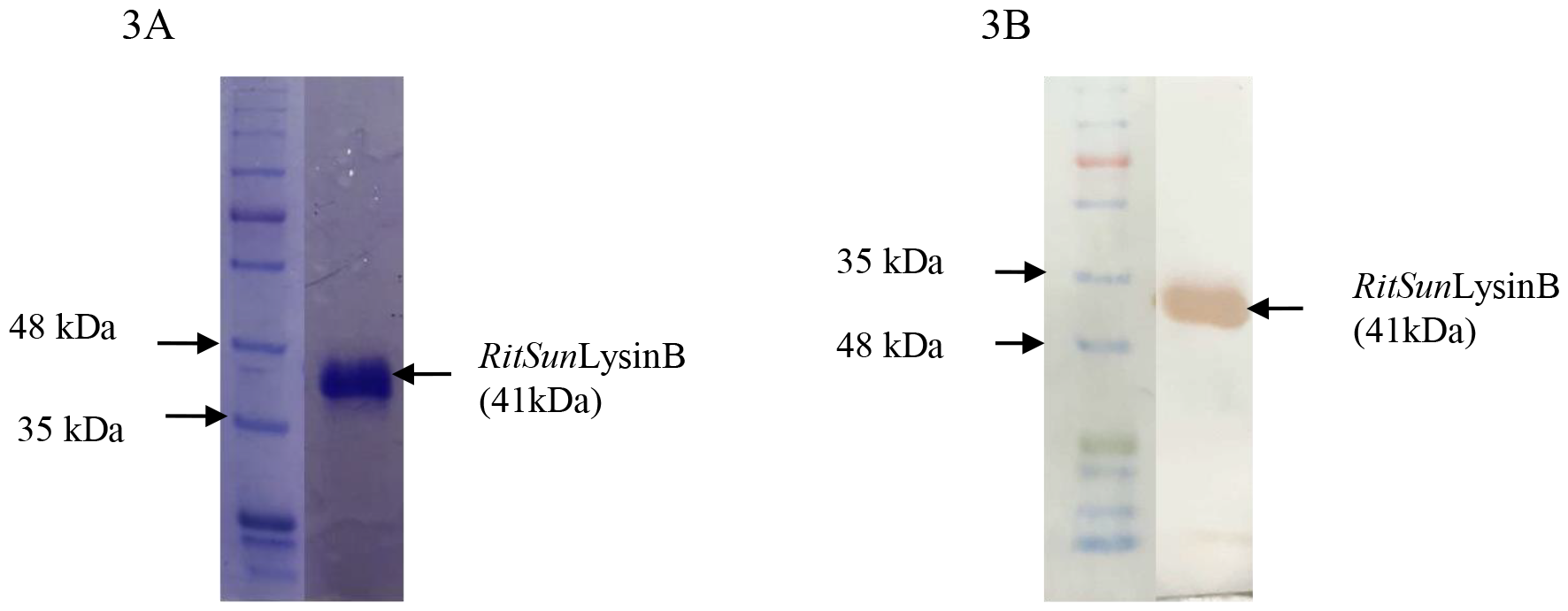
His-tagged purified recombinant *RitSun* LysinB (41 kDa). **3A**. SDS-PAGE (12%), **3B**. Western Blot

### 3. *In vitro* activity assay (pNPB assay) for measuring LysinB activity

The esterase activity of LysinB was estimated at different enzyme concentrations and different temperatures, using pNPB as the substrate. The release of pNP was measured at 410 nm (Figure 4A). The enzyme activity was stable up to 55°C (Figure 4B). The specific activity of *RitSun* LysinB and LysinB reported from other mycobacteriophages is shown in Table 1.

**Figure 4.**
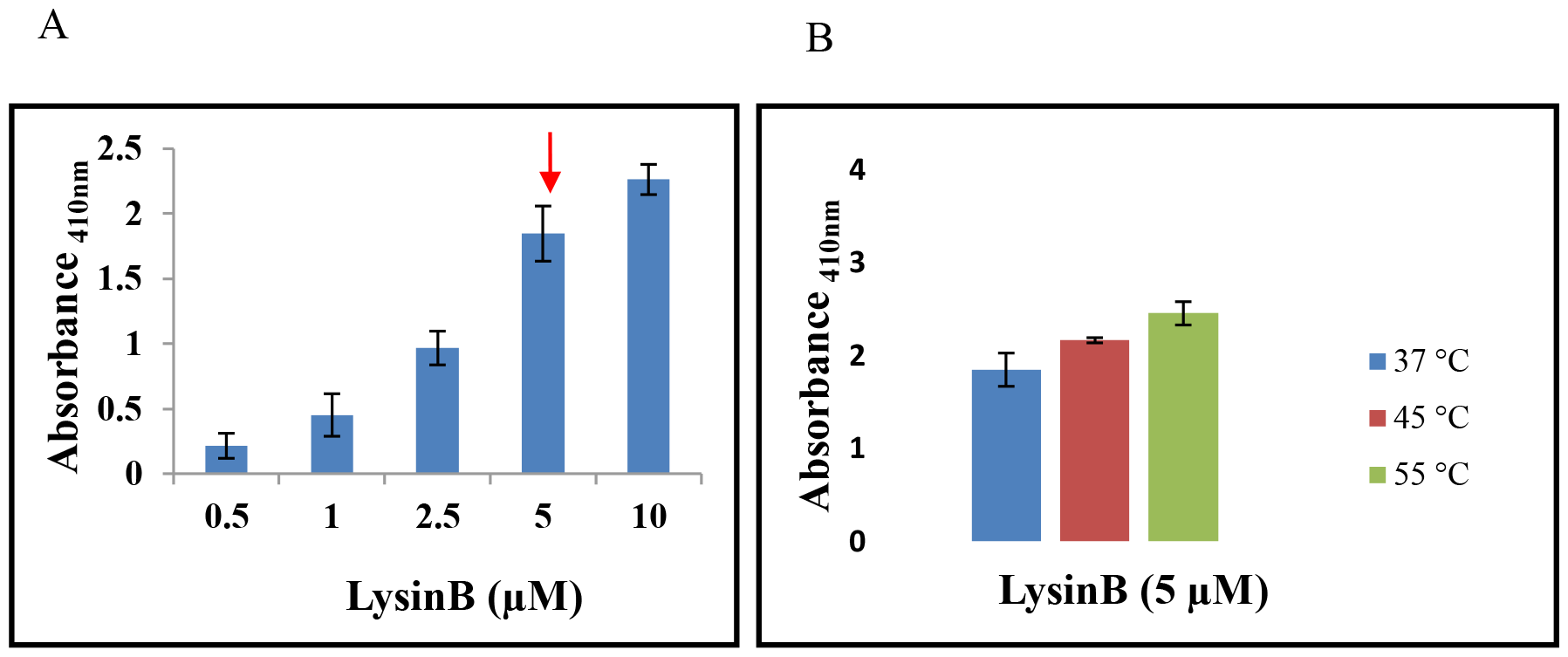
**A**. Esterase assay of *RitSun* LysinB at different concentrations for an incubation period of 15 minutes at 37°C. The substrate concentration of pNPB was 10 mM (in 25 mM Tris buffer, pH 7.2). The enzyme reaction was performed twice in duplicate sets. **Figure 4B**. Esterase assay to study the effect of temperature on *RitSun* LysinB activity. The concentration of pNPB was 10 mM in (25 mM Tris buffer, pH 7.2) and the absorbance was measured at 410 nm. The enzyme reaction was performed twice in duplicate sets.

### 4. Antibacterial effect of *RitSun* LysinB on *M. smegmatis* Mc^2^ 155

#### 4.1 Spot Test, Turbidity Reduction Method (TRM) and Growth Inhibitory Effect

*RitSun* LysinB spot test (in the presence of 0.05% Tween-80) showed a zone of clearance on *M. smegmatis* lawn after 48 hours of incubation at 37°C (Figure 5A), demonstrating its cell lytic activity from without. The TRM analysis (Figure 5B) showed a 45.6%±2.5% reduction in *M. smegmatis* Mc^2^ 155 growth after 24 hours of treatment (15 μM). In untreated cells, the increase in growth after 24 hours was 232.1%±9.5% compared with the growth at zero hours. To test the growth inhibitory effect of *RitSun* LysinB, dilutions of enzyme-treated *M. smegmatis* cells were spotted on 7H10 plates. At 10^−2^ dilution of the LysinB (15 μM) treated cells, no bacterial colonies were observed after 24 hours of incubation (Figure 5C).

**Figure 5A.**
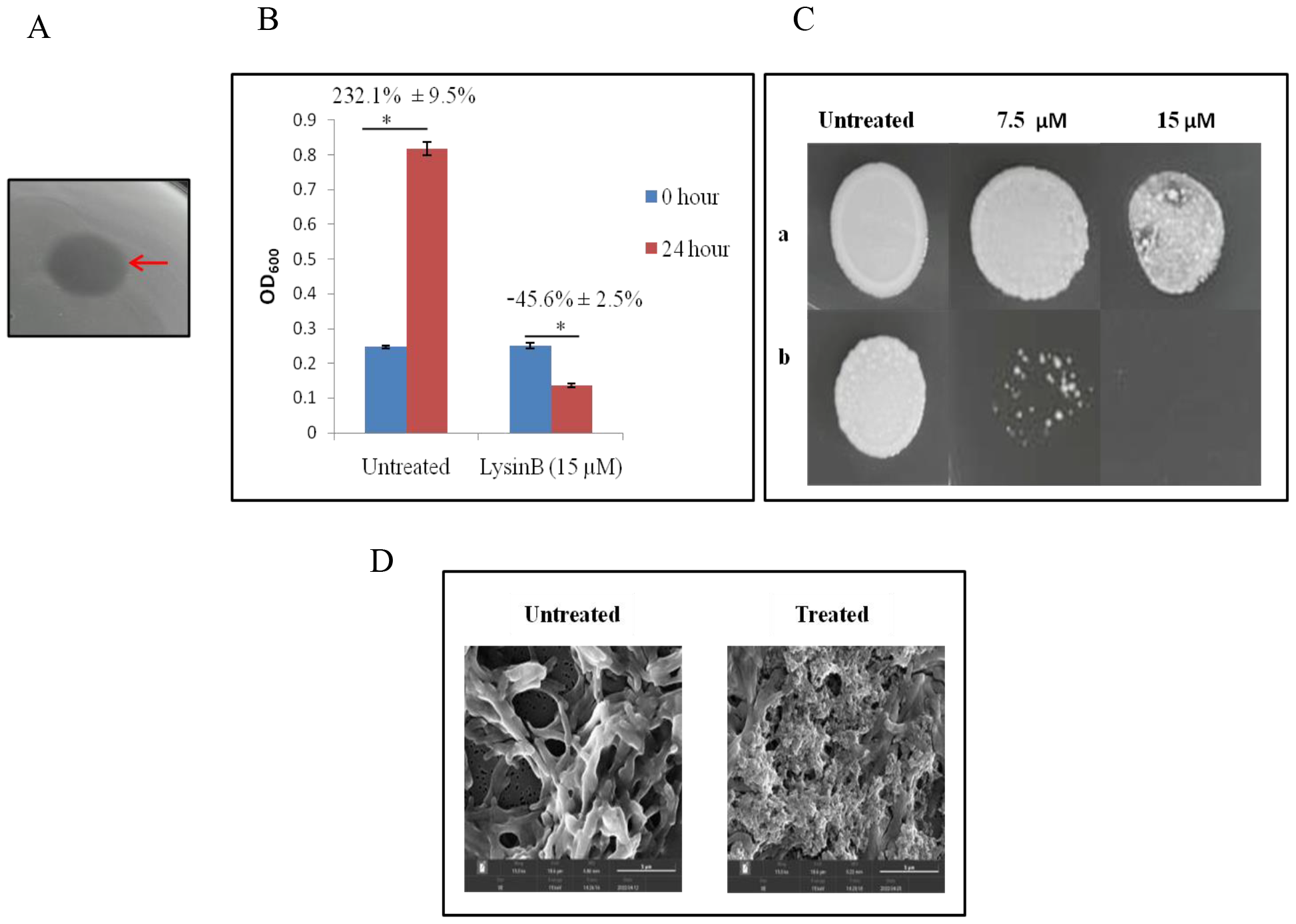
Spot test of *RitSun* LysinB on *M. smegmatis* growth in Tween-80 (0.05%) containing 7H10 media. **Figure 5B**. Turbidity Reduction Method (TRM). Antimycobacterial activity of *RitSun* LysinB (measured at OD_600_) on *M. smegmatis* Mc^2^ 155 at 0 and 24 hours of treatment.

#### 4.2 FESEM analysis of *RitSun* LysinB treated planktonic cells

To observe the effect of the enzyme on cell morphology, *M. smegmatis* cells were treated with LysinB (7.5 μM) for 4 hours at 37°C and analysed by Field Emission Scanning Electron Microscopy (FESEM) (Figure 5D). Compared to the untreated cells, the morphology of *RitSun* LysinB**-**treated cells was disrupted, indicating the cell-damaging effect of LysinB. Structural damage to the cell wall can affect the integrity of bacterial cells and result in cell death [10]. LysinB targets mycobacteria due to its activity as a mycolylarabinogalactan esterase, which hydrolyzes the ester bond that connects the mycolic acid-rich outer membrane to arabinogalactan. This LysinB capacity most likely mechanically weakens and disrupts the bacterial cell wall structure and causes disruption in the cell wall [29]. The ability of our endolysin to cause mycobacterial cell (*M. smegmatis*) lysis on external application, without the aid of a permeabilizing agent, is significant and can find clinical applications.

The percentage decrease in growth was calculated by taking growth at 0 hours as 100%. The enzyme concentration was 15 μM, and the experiment was performed twice in duplicate sets. **Figure 5C**. Growth inhibitory effect of different concentrations of *RitSun* LysinB on *M. smegmatis* Mc^2^ 155 after 24 hours of incubation at 37°C. 1-Untreated cells; 2-Treatment with 7.5μM *RitSun* LysinB; 3-Treatment with 15μM *RitSun* LysinB: ‘a’ represents growth at dilution 10^−1^ and ‘b’ at 10^−2^. **Figure 5D**. FESEM images of *M. smegmatis* Mc^2^ 155 cells treated with *RitSun* LysinB (7.5 μM) for 4 hours at 37°C. The images were acquired at 15 Kx magnification. The assays were performed twice in duplicate sets.

### 5. Inhibitory effect of *RitSun* LysinB on *M. smegmatis* Mc^2^ 155 biofilm formation

Biofilms are notorious for causing recalcitrant infections and are also responsible for drug resistance. Unlike several antibiotics, the ability of endolysins to kill both metabolically active and inactive bacterial cells and target biofilms makes them more versatile [36]. We used the Crystal Violet staining method to assess the impact of *RitSun* LysinB on *M. smegmatis* biofilm formation and noted a strong inhibitory effect of 79.18%±2.20% (Figure 6). The anti-biofilm effect is important for clinical applications and further supports the efficacy of this novel enzyme, especially since the role of endolysins on mycobacterial biofilm formation hasn’t yet been reported.

**Figure 6.**
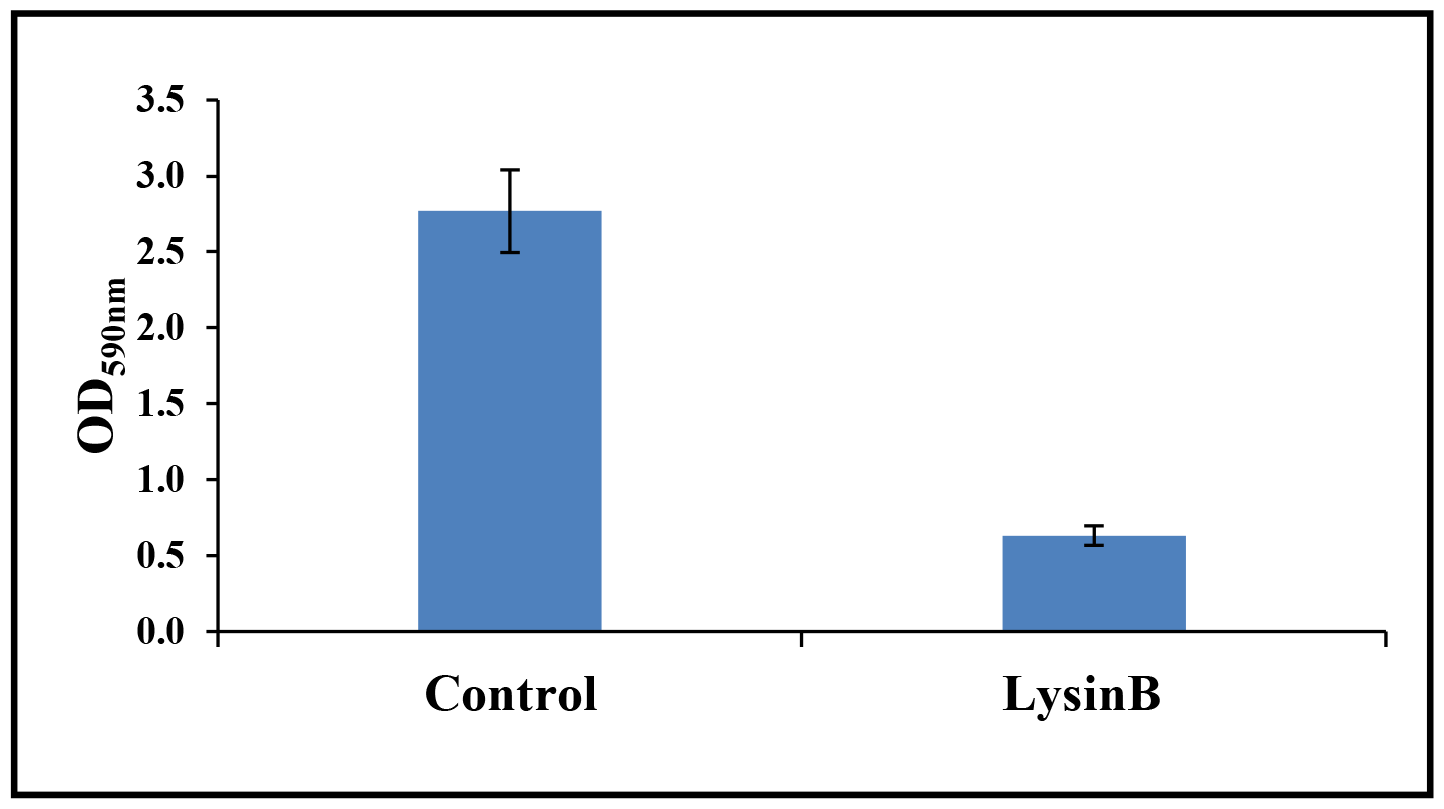
Inhibitory effect of *RitSun* LysinB on *M. smegmatis* Mc^2^ 155 biofilm formation under static conditions. Control: *M. smegmatis* Mc^2^155 biofilm formation after 96 hours incubation at 37°C; Treatment group: LysinB added to *M. smegmatis* Mc^2^ 155 before biofilm formation.The assay was performed twice in duplicate sets.

## CONCLUSION

Endolysins are synthesized in the latter stages of the phage infection cycle. They impair the host’s cell wall by damaging the peptidoglycan, allowing the progeny to release and infect the neighbouring bacterial cells. The characterization and administration of purified recombinant lytic enzymes have been effectively tested over the last several years in different animal models of human illnesses [29,37,38,39,40], and their commercial use has also been approved in some cases [21,41].

The intricate structure of the mycobacterial cell envelope sets it apart from the other bacteria. Mycobacteria have peptidoglycan covalently bonded to an arabinogalactan polymer, which is then esterified to mycolic acids known as the mycolyl arabinogalactan peptidoglycan (mAGP) complex. Mostof the mycobacteriophage genomes encode for a second Lysin, known as LysinB, which acts upon the mAGP complex. The LysinB enzymes have an esterase activity that breaks the ester linkage between mycolic acid and arabinogalactan layer and hence is being considered a more compelling candidate for therapeutic protein than LysinA.

In this work, we have performed structural (*in silico)* and functional (*in vitro*) analysis of the predicted LysinB from *RitSun* mycobacteriophage, which belongs to the sub-cluster F2 and was isolated from a soil sample near a hospital in Delhi. Compared with the previously reported LysinB from other phages, namely, *D29, Ms6* and *PDRPxv*, we found the specific activity (esterase) of *RitSun* LysinB to be the highest. Also, its intrinsic property to permeate through *M. smegmatis* cell wall to cause cell lysis and the anti-biofilm activity indicates the enzyme’s potential as a promising candidate for developing antimycobacterial enzybiotics. Though we found *RitSun* LysinB specific to *M. smegmatis*, further studies to identify the sequence with the permeabilizing activity could help develop fusion lysins that are effective against the pathogenic species of mycobacteria.

## MATERIALS AND METHODS

### *In silico* characterization of LysinB

#### Analysis of *RitSun* LysinB sequence

The LysinB gene sequence was identified from the *RitSun* mycobacteriophage genome (GenBank ID:MZ343158.1). The phylogenetic analysis of *RitSun* LysinB was performed using VipTree software [42]. The protein sequence was analysed using BLASTp [43], InterProScan [44] and HMMER [45] based on sequence homology and domain organisation. The Clustal Omega tool [46] was used for multiple sequence alignment. A comparative analysis of *RitSun* LysinB with sequences from other F sub-clusters phages was performed using BLASTp and InterProScan.

#### Cloning and Expression of LysinB

The LysinB genes-specific primers for PCR amplification (LysBF: 5’-CCG<u>GAATTC</u>ATGAAACTCAACGGGGTATATG-3’); LysBR: 5’-CCC<u>AAGCTT</u>TCATGCGACACCCCTCATC-3’) were synthesised by Eurofins, India and pET28a (Novagen, USA), a gift from Dr. S. Ramachandran (Institute of Genomics and Integrative Biology, New Delhi) was used as a cloning vector.

The cloned gene was expressed in *E*.*coli* BL21 (DE3), as previously described by *Eniyan et al*. [13]. *E*.*coli* BL21 (DE3) cells transformed with pET28a-LysinB plasmid were grown in LB media to an OD_600_ of 0.4-0.6. The induction parameters for the overexpression of the LysinB gene were optimized at varying concentrations of IPTG and different incubation periods and temperatures. The final *RitSun* LysinB induction conditions were achieved using 1mM IPTG and incubation at 37ºC for 3 hours at 200 rpm. After induction, the cells were harvested and resuspended in lysis buffer (25 mM Tris, pH 7.9; 0.5 M NaCl, 10 mM Imidazole, 1 mM PMSF). The cell suspension was sonicated at 90% amplitude with 30 sec each on/off pulse for ten cycles, followed by centrifugation of the lysate at 13,000 rpm,4ºC for 45 minutes. The soluble and pellet fractions were analysed for the expression of the recombinant LysinB.

#### Purification of *RitSun* LysinB

The recombinant endolysin protein was purified using Ni-NTA affinity chromatography, as described previously [13]. Briefly, soluble fraction and Ni-NTA matrix agarose (Qiagen, Germany) were kept for binding for two hours at 4ºC. The agarose gel bound with the His-tagged recombinant protein was packed in the chromatography column (Qiagen, Germany), and the flow through was discarded. The column was washed twice with increasing concentrations of imidazole (20 & 40 mM), and the protein was eluted at 250 mM imidazole. The eluted fractions were analyzed by SDS-PAGE (12 %) and dialysed in a buffer containing 25 mM Tris and 150 mM NaCl.

#### Western Blotting

The purified protein was transferred from the gel to the PVDF membrane for 2 hours at 90V. The membrane was blocked with 3% BSA at 4ºC and incubated with 1:2000 dilution of mouse anti-His antisera (Santacruz Biotech, USA) for 4 hours. Post-incubation, the membrane was washed once with 1X PBS, thrice with 1X PBS-T (containing 0.05% Tween-80) and again with 1X PBS and incubated with 1:10,000 anti-mouse IgG HRP-conjugated antibody (Santacruz Biotech, USA) for 45 minutes and washed as mentioned above. The membrane blot was developed using 3,3-diaminobenzidine (DAB) as the chromophore and H_2_O_2_ as the substrate [13,47].

#### *In vitro* assay of *RitSun* LysinB

The esterase activity of the LysinB enzyme was estimated by p-nitrophenol butyrate (pNPB) assay [29]. Release of p-Nitrophenol (PNP) was measured by incubating different concentrations (0.5-10 μM range) of LysinB with pNPB (10 mM in 25 mM Tris buffer, pH 7.2) at 37°C for 15 minutes, and the absorbance was measured at 410 nm. Each experiment was performed twice independently (in duplicates), and the results were calculated as average ± SD.

#### Antibacterial Effect of LysinB using plate lysis and Turbidity Reduction Method (TRM)

LysinB protein was spotted on *M. smegmatis* Mc^2^ 155 lawn on 7H10 plates and incubated at 37ºC until a clearance zone appeared. For microtiter plate-based TRM, *M. smegmatis* Mc^2^ 155 cells, adjusted to an OD_600_ to 0.3, were treated with *RitSun* LysinB enzyme (15 μM)andincubated at 37°C for 24 hours under shaking conditions (200 rpm). OD_600_ was measured at 0 hours and after 24 hours of incubation. P-values were calculated to obtain a level of significance for each treatment group. The bacterial culture without adding the enzyme was treated as a control. The treated and the untreated cells were serially diluted and spotted on 7H10 plates containing tween-80 (0.05%) and incubated at 37°C for 48 hours. Each experiment was performed twice independently (in duplicates).

### Field Emission Scanned Electron Microscopy (FESEM) of LysinB-treated *M. smegmatis* cells

For the FESEM analysis, the protocol by *Lai et al*. [10] was followed. *M. smegmatis* Mc^2^ 155 planktonic culture was grown in 7H9 complete media and Lysin B was added at <15 μM concentrations. Aliquots (500μl) of *M. smegmatis* Mc^2^ 155 cells adjusted to an OD_600_ of 0.5 were treated with LysinB and incubated at 37°C for 4 hours. Treated cells were centrifuged at 10,000 rpm for 5 minutes, and the cell pellet was washed with 1X PBS. The washed cells were fixed overnight in 2.5% glutaraldehyde at 4°C. Glutaraldehyde and the fixed cells were washed once with 1X PBS, followed by gradient dehydration with ethanol (5 min each in 60%, 80%, 90% and 10 min in 100% ethanol). The dehydrated cells were resuspended in 1X PBS+Tween-80 (0.05%) and placed on a 0.2μm Polycarbonate (PC) membrane. The membrane was air-dried and mounted on a stub for gold sputter coating. Samples were visualized in TESCAN (at the Indian Institute of Technology, New Delhi) at 15 kX magnification.

#### Anti-biofilm activity of LysinB

To examine the inhibitory effect of *RitSun* LysinB on *M. smegmatis* Mc^2^ 155 biofilm formation, the *Kiefer et al*. [48] protocol was followed with minor modifications. Briefly, *M. smegmatis* cells (OD_600_ adjusted to 0.1) were mixed with LysinB in a 24-well plate and incubated at 37°C for 96 hours. Biofilms were quantified using the crystal violet (CV) staining method. The experiments were performed thrice independently (in triplicates each time), and the results were calculated as an average of the percentage inhibition ± SD observed over untreated biofilm.

#### Crystal Violet Assay

After completing LysinB treatment, the planktonic cells were aspirated, and the biofilm was air-dried and stained with freshly prepared crystal violet stain (1%) for 30 minutes at 37°C and washed twice with 1X PBS. The CV stain was extracted with 95% ethanol, and the absorbance was measured at 590 nm [48].

## ACKNOWLEDGEMENTS

We thank the Science and Engineering Research Board (SERB), New Delhi, India, for supporting the project (EMR/2017/004051). RA received the Senior Research Fellowship (SRF) from University Grants Commission (UGC), New Delhi, India. KN received the Innovation in Science Pursuit for Inspired Research (INSPIRE) fellowship from the Department of Science and Technology (DST), India. We thank Acharya Narendra Dev College (ANDC), University of Delhi, New Delhi, India, for providing the infrastructural support.

## AUTHOR CONTRIBUTIONS

UB conceptualized, designed, and supervised the experiments and edited the manuscript. RA performed the experiments, prepared and edited the original draft. KN performed the Western Blotting and the Biofilm experiment. All the authors have read and approved the final manuscript.

## DATA AVAILABILITY STATEMENT

All the data generated and analyzed during this study is included in this published article or its supplementary files.

## REFERENCES

1. Global TB report, 2022.

2. Dadgostar, P. Antimicrobial resistance: implications and costs. Infection and drug resistance, 3903–3910 (2019).

3. Majumder, M. A. A. et al. Antimicrobial stewardship: Fighting antimicrobial resistance and protecting global public health. Infection and drug resistance, 4713–4738 (2020).

4. Allué-Guardia, A., Saranathan, R., Chan, J., & Torrelles, J. B. Mycobacteriophages as potential therapeutic agents against drug-resistant tuberculosis. International Journal of molecular sciences, 22(2), 735 (2021).

5. Zulu, M., Monde, N., Nkhoma, P., Malama, S., & Munyeme, M. Nontuberculous mycobacteria in humans, animals, and water in Zambia: A systematic review. Frontiers in Tropical Diseases, 2, 9 (2021).

6. Tarashi, S., Siadat, S. D., & Fateh, A. Nontuberculous mycobacterial resistance to antibiotics and disinfectants: Challenges still ahead. BioMed Research International, (2022).

7. Esteves, N. C., & Scharf, B. E. Flagellotropic bacteriophages: opportunities and challenges for antimicrobial applications. International Journal of Molecular Sciences, 23(13), 7084 (2022).

8. Schmelcher, M., Donovan, D. M., & Loessner, M. J. Bacteriophage endolysins as novel antimicrobials. Future microbiology, 7(10), 1147–1171 (2012).

9. Abdelrahman, F. et al. Phage-encoded endolysins. Antibiotics, 10(2), 124 (2021).

10. Lai, M. J. et al. Antimycobacterial activities of endolysins derived from a mycobacteriophage, BTCU-1. Molecules, 20(10), 19277–19290 (2015).

11. Gerstmans, H., Rodríguez-Rubio, L., Lavigne, R., & Briers, Y. From endolysins to Artilysin® s: novel enzyme-based approaches to kill drug-resistant bacteria. Biochemical Society Transactions, 44(1), 123–128 (2016).

12. Love, M. J., Bhandari, D., Dobson, R. C., & Billington, C. Potential for bacteriophage endolysins to supplement or replace antibiotics in food production and clinical care. Antibiotics, 7(1), 17 (2018).

13. Eniyan, K., Sinha, A., Ahmad, S., & Bajpai, U. Functional characterization of the endolysins derived from mycobacteriophage PDRPxv. World Journal of Microbiology and Biotechnology, 36, 1–11 (2020).

14. Payne, K., Sun, Q., Sacchettini, J., & Hatfull, G. F. Mycobacteriophage Lysin B is a novel mycolylarabinogalactan esterase. Molecular microbiology, 73(3), 367–381 (2009).

15. Catalão, M. J., & Pimentel, M. Mycobacteriophage lysis enzymes: targeting the mycobacterial cell envelope. Viruses, 10(8), 428 (2018).

16. Zermeño-Cervantes, L. A., Martínez-Díaz, S. F., Venancio-Landeros, A. A., & Cardona-Félix, C. S. Evaluating the efficacy of endolysins and membrane permeabilizers against Vibrio parahaemolyticus in marine conditions. Research in Microbiology, 174(7), 104104 (2023).

17. Gontijo, M. T. P., Jorge, G. P., & Brocchi, M. Current status of endolysin-based treatments against Gram-negative bacteria. Antibiotics, 10(10), 1143 (2021).

18. Rahman, M. U., et al. Endolysin, a promising solution against antimicrobial resistance. Antibiotics, 10(11), 1277 (2021).

19. Vazquez, R., Garcia, E., & Garcia, P. Phage lysins for fighting bacterial respiratory infections: a new generation of antimicrobials. Front Immunol 9: 2252 (2018).

20. Nelson, D., Loomis, L., & Fischetti, V. A. Prevention and elimination of upper respiratory colonization of mice by group A streptococci by using a bacteriophage lytic enzyme. Proceedings of the National Academy of Sciences, 98(7), 4107–4112 (2001).

21. Murray, E., Draper, L. A., Ross, R. P., & Hill, C. The advantages and challenges of using endolysins in a clinical setting. Viruses, 13(4), 680 (2021).

22. Watson, A., et al. Antimicrobial activity of exebacase (lysin CF-301) against the most common causes of infective endocarditis. Antimicrobial Agents and Chemotherapy, 63(10), 10–1128 (2019).

23. Jun, S. Y. et al. Pharmacokinetics and tolerance of the phage endolysin-based candidate drug SAL200 after a single intravenous administration among healthy volunteers. Antimicrobial agents and chemotherapy, 61(6), 10–1128 (2017).

24. Totté, J. E., van Doorn, M. B., & Pasmans, S. G. Successful treatment of chronic Staphylococcus aureus-related dermatoses with the topical endolysin Staphefekt SA. 100: a report of 3 cases. Case reports in dermatology, 9(2), 19–25 (2017).

25. Dedrick, R. M. et al. Engineered bacteriophages for treatment of a patient with a disseminated drug-resistant Mycobacterium abscessus. Nature medicine, 25(5), 730–733 (2019).

26. Dedrick, R. M. et al. Phage therapy of Mycobacterium infections: compassionate use of phages in 20 patients with drug-resistant mycobacterial disease. Clinical infectious diseases, 76(1), 103–112 (2023).

27. Hosseiniporgham, S., & Sechi, L. A. A Review on Mycobacteriophages: From Classification to Applications. Pathogens, 11(7), 777 (2022).

28. Davies, C. G., Reilly, K., Altermann, E., & Hendrickson, H. L. PLAN-M; Mycobacteriophage Endolysins Fused to Biodegradable Nanobeads Mitigate Mycobacterial Growth in Liquid and on Surfaces. Frontiers in Microbiology, 12, 562748 (2021).

29. Fraga, A. G. et al. Antimicrobial activity of Mycobacteriophage D29 Lysin B during Mycobacterium ulcerans infection. PLoS neglected tropical diseases, 13(8), e0007113 (2019).

30. Singh, A. K., et al. Mycobacteriophage D29 Lysin B exhibits promising anti-mycobacterial activity against drug-resistant Mycobacterium tuberculosis. Microbiology Spectrum, 11(6), e04597–22 (2023).

31. Griego, A. et al. Endolysin B: A new archetype in M. tuberculosis treatment. bioRxiv, 2023–12 (2023).

32. Korany, A. H. et al. Comparative structural analysis of different mycobacteriophage-derived mycolylarabinogalactanesterases (Lysin B). Biomolecules, 10(1), 45 (2019).

33. Gil, F., et al. Mycobacteriophage Ms6 LysB specifically targets the outer membrane of Mycobacterium smegmatis. Microbiology, 156 (Pt 5), 1497 (2010).

34. phagesdb.org

35. Abouhmad, A., Korany, A. H., Grey, C., Dishisha, T., & Hatti-Kaul, R. Exploring the enzymatic and antibacterial activities of novel mycobacteriophage lysin B enzymes. International Journal of Molecular Sciences, 21(9), 3176 (2020).

36. Fursov, M. V. et al. Antibiofilm activity of a broad-range recombinant endolysin LysECD7: in vitro and in vivo study. Viruses, 12(5), 545 (2020).

37. Loeffler, J. M., & Fischetti, V. Synergistic lethal effect of a combination of phage lytic enzymes with different activities on penicillin-sensitive and-resistant Streptococcus pneumoniae strains. Antimicrobial agents and chemotherapy, 47(1), 375–377 (2003).

38. Cheng, Q., Nelson, D., Zhu, S., & Fischetti, V. A. Removal of group B streptococci colonizing the vagina and oropharynx of mice with a bacteriophage lytic enzyme. Antimicrobial agents and chemotherapy, 49(1), 111–117 (2005).

39. Fischetti, V. A., Nelson, D., & Schuch, R. Reinventing phage therapy: are the parts greater than the sum?. Nature biotechnology, 24(12), 1508–1511 (2006).

40. Grandgirard, D., Loeffler, J. M., Fischetti, V. A., & Leib, S. L. Phage lytic enzyme Cpl-1 for antibacterial therapy in experimental pneumococcal meningitis. The Journal of infectious diseases, 197(11), 1519–1522 (2008).

41. Gutierrez, D., Fernandez, L., Rodriguez, A., & Garcia, P. Are phage lytic proteins the secret weapon to kill Staphylococcus aureus? mBio 9: e01923–17 (2018).

42. Nishimura, Y., et al. ViPTree: the viral proteomic tree server. Bioinformatics, 33(15), 2379–2380 (2017).

43. Altschul, S. F., Gish, W., Miller, W., Myers, E. W., & Lipman, D. J. Basic local alignment search tool. Journal of molecular biology, 215(3), 403–410 (1990).

44. Jones, P. et al. InterProScan 5: genome-scale protein function classification. Bioinformatics, 30(9), 1236–1240 (2014).

45. Eddy, S. R. Profile hidden Markov models. Bioinformatics (Oxford, England), 14(9), 755–763 (1998).

46. Madeira, F. et al. The EMBL-EBI search and sequence analysis tools APIs in 2019. Nucleic acids research, 47(W1), W636–W641 (2019).

47. Mahmood, T., & Yang, P. C. Western blot: technique, theory, and trouble shooting. North American Journal of medical sciences, 4(9), 429 (2012).

48. Kiefer, B., & Dahl, J. L. Disruption of Mycobacterium smegmatis biofilms using bacteriophages alone or in combination with mechanical stress. Advances in Microbiology, 5(10), 699 (2015).

